# Spontaneous Emergence of Symmetry in a Generative Model of Protein Structure

**DOI:** 10.1101/2025.11.03.686219

**Authors:** Anton Oresten, Kenta Sato, Aron Stålmarck, Lukas Billera, Hedwig Nora Nordlinder, Jack Collier Ryder, Mateusz Kaduk, Ben Murrell

**Affiliations:** Department of Microbiology, Tumor and Cell Biology, Karolinska Institutet, Stockholm, Sweden

**Keywords:** Protein design, Flow matching

## Abstract

Generative models are becoming powerful tools for protein design, enabling the creation of novel protein structures and sequences. Recent approaches have shown success using diffusion models and flow matching to sample realistic protein folds. Many methods explicitly incorporate structural biases or constraints—for example, enforcing symmetry during generation—to steer the design process. Here we report the spontaneous emergence of structural symmetry in a transformer-based generative model for proteins, without any symmetry-specific conditioning during training or constraints during generation. In our flow-matching model, a single attention head in an SE(3)-equivariant transformer layer was found to be primarily responsible for the model’s ability to generate symmetric arrangements of backbone residues across chains, or repeating motifs within a chain. Our results show that protein generative models can learn high-level structural patterns implicitly from training data. This opens new questions about interpretability and control in generative design: understanding how and why a single attention head can govern a complex global property like symmetry may inform future model architectures and help exploit emergent behaviors for better protein engineering.

## Introductionss

Deep generative modeling has begun to reshape protein engineering, providing a statistical complement to physics-based approaches [1], [2]. These include i) “deep network hallucination” approaches that propagate gradients through a pre-trained folding model to search for a sequence that folds into the desired target structure [3], ii) a class of related diffusion and flow matching models that learn to transport an easy-to-sample-from distribution to the distribution of proteins, and iii) autoregressive backbone models that sample the placement of each successive backbone amino acid, one after another [4].

Diffusion-style generators dominate current benchmarks, but continuous normalizing-flow variants— trained by **flow matching** [5]—offer exact likelihoods and fast sampling, and have already been adapted to SE(3)-equivariant protein backbones [6] and to joint sequence–structure “co-design” on discrete state-spaces [7].

Here we train a multimer-aware transformer whose backbone flow evolves via Brownian bridges on SO(3) and ℝ^3^ and whose discrete sequence component is learned via Discrete Flow Matching [7], [8]. The architecture inherits Invariant Point Attention from AlphaFold2 [9] and employs self-conditioning to stabilize long trajectories [10]. Crucially, the training objective contains no symmetry term, and the initial noise field places each chain independently.

## Results

As described in Methods, ChainStorm uses flow matching [5], where a model is trained to transport a simple distribution that is easy to sample from to a complex distribution which is, in our case, the protein structures in the Protein Data Bank (PDB) [11].

ChainStorm is trained to generate protein backbone and sequence jointly, and the model takes a noise-corrupted backbone, where the backbone atoms of amino acid residues are represented as rigid “frames”, each with an SO(3) orientation and ℝ^3^ position (following [6], [7], [9]), and a noised discrete amino acid sequence, which is a number between 1 and 21 (where 21 is a “dummy” amino acid generated by the noising process).

ChainStorm uses an Invariant Point Attention (IPA) [9] transformer-based architecture with a central role for self-conditioning [10], where the model’s previous output is included as input for the next step. As shown in Figure 1, each layer in the model (of which there are six) first allows communication between the selfconditioning frames, then between the self-conditioning frames and the current frames, and then finally between the current frames, after which the position of the current frames is updated.

**Figure 1:**
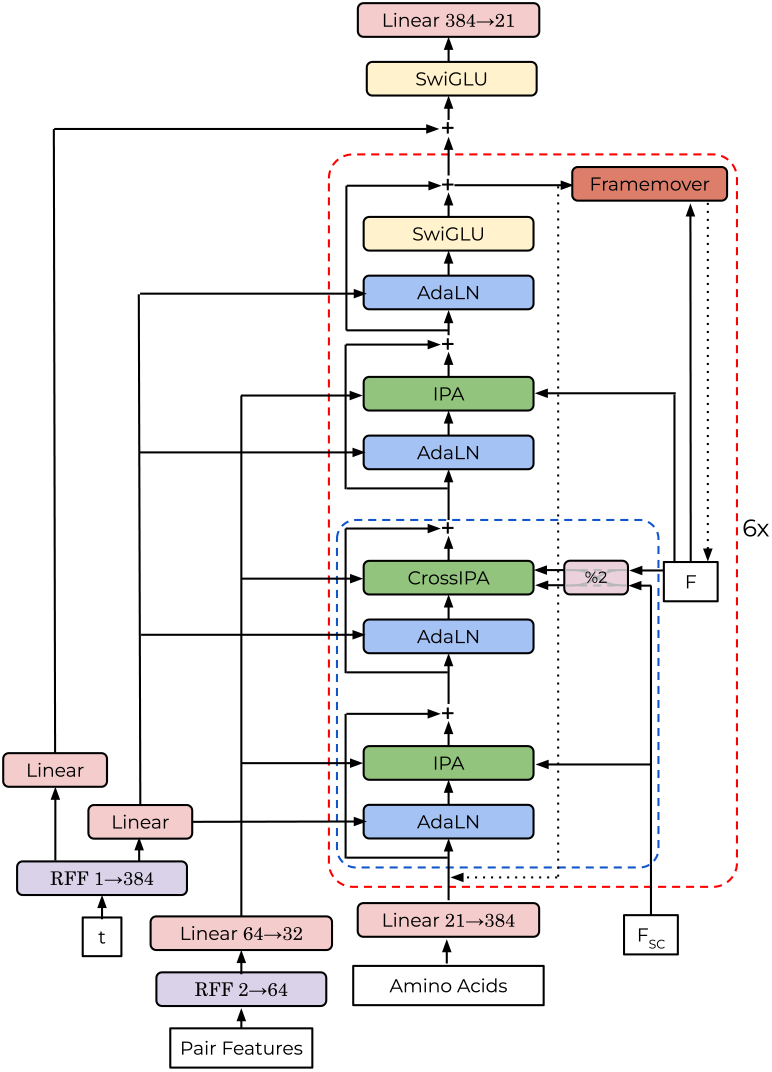
Model architecture. The model takes the diffusion time *t*, the current backbone “frames”, *F*, the current Amino Acids (onehot encoded), and a set of Pair Features that encode the distance between two residues in the primary sequence, and whether or not two residues are part of the same chain. The model also optionally takes *F*_*SC*_, which are the predicted backbone frames from the previous pass of the model. The initial residue embeddings are projected from the amino acids, and updated by the model “trunk” which repeats 6 times, and comprises IPA self-attention on *F*_*SC*_, IPA crossattention between *F*_*SC*_ and *F* (where the frames that are used for keys vs queries alternates each layer), and IPA self-attention on *F*. The pair features are encoded via Random Fourier Features (RFF) and fed into each IPA layer, and diffusion time *t* is encoded via RFF and fed to an adaptive layer normalization (AdaLN) prior to each IPA layer, as well as prior to the SwiGLU feedforward layer. In each iteration of the trunk, the frames *F* are updated via a function of a linear projection of the embeddings (as in AlphaFold2). The model outputs the final state of *F*, and the amino acid logits which are projected from a SwiGLU feedforward layer after completion of the trunk.

We train on multi-chain structures (see Data Preparation), and, in addition to the frames, each IPA layer is provided with an encoding of relative position of all pairs of amino acids in the primary sequence, as well as whether or not any two amino acids are from the same protein chain.

We use a “multimodal” flow matching approach similar to Campbell et al [7] in that the backbone and sequence are modeled jointly, but where ChainStorm’s backbone flow involves stochastic bridges (similar to FoldFlow SFM in [6]) rather than deterministic geodesic paths. Examples of our conditional paths are depicted in Figure 9.

The generative process begins with randomly placed frames, with uniformly distributed orientations, unit-variance (in nanometers) Gaussian positions, and dummy 21st amino acid states. The only user-controllable inputs are the residue numbers and the chain IDs. If the residue numbers are contiguous (i.e. no missing structure), then the only user input is the length of each chain to be generated.

### Structure emerges early during generation

To understand the timecourse of the generative process, focusing on the backbone generation, we visualizein stances of the “current” frames *X*_*t*_, with the model’s corresponding prediction of its final state 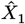. As can be seen in Figure 2, very early in the generative process, typically between *t* = 0 and *t* = 0.1 the model has a clear prediction of the macroscopic details of its own terminal state.

**Figure 2:**
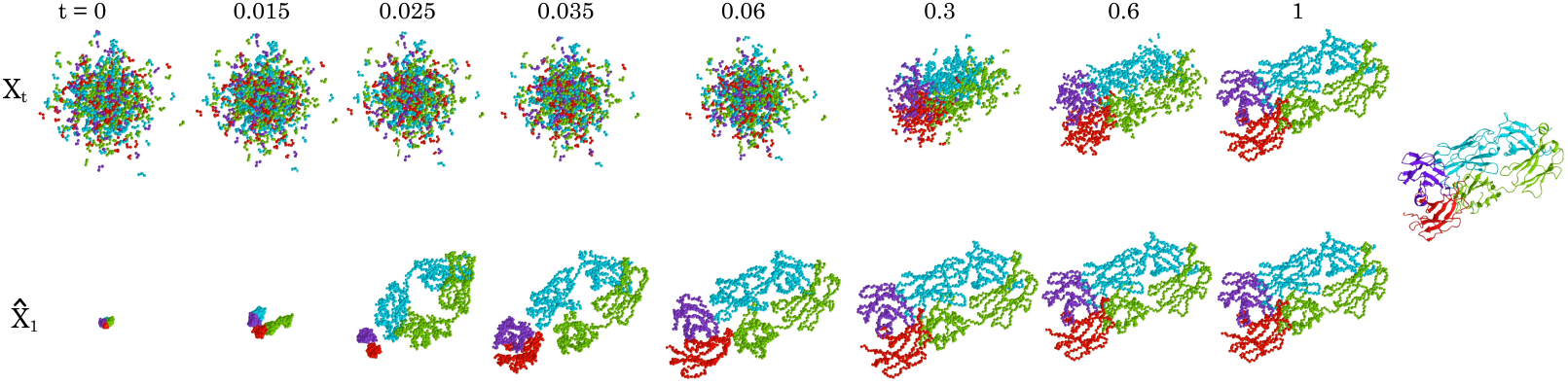
Inference trajectory visualization. The generative process is shown, from left to right, for *t* from 0 to 1 (note: not uniformly spaced). The top row shows *X*_*t*_, which tracks the current backbone frames, and the bottom shows 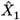, the model’s current prediction of its final state. While *X*_*t*_ is still indistinguishable from a Gaussian point cloud, by *t* = 0.06 the macroscopic details of the final structure (ribbon diagram, bottom) are already clear in 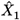. Also note how the antibody FAB-like dimer emerges slightly ahead of the other two chains, a pattern commonly observed for immunoglobulin domains.

Since ChainStorm employs no coupling between *X*_0_ and *X*_1_ (i.e. no rectified flow or optimal transport), the initial 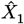 prediction at *t* = 0 lies in a tight ball at the center of the *X*_0_ frame cloud. The first event is the separation of the different chains from each other, which governs their final placement, followed quickly by the layout of the chains themselves, and then proceeding with progressively more minor backbone adjustments as *t* → 1.

### ChainStorm consistently generates symmetric structures

Despite not employing any symmetry constraints during generation, nor being explicitly provided with symmetry information during training, a visual inspection of the output suggested that ChainStorm spontaneously generates structures that exhibit, at least approximately, symmetry. Figure 3 shows examples of dimers of heterodimers, which are reliably generated by the model when generating four chains and the chain lengths (which are the only model input) are consistent with a dimer of heterodimer arrangement.

**Figure 3:**
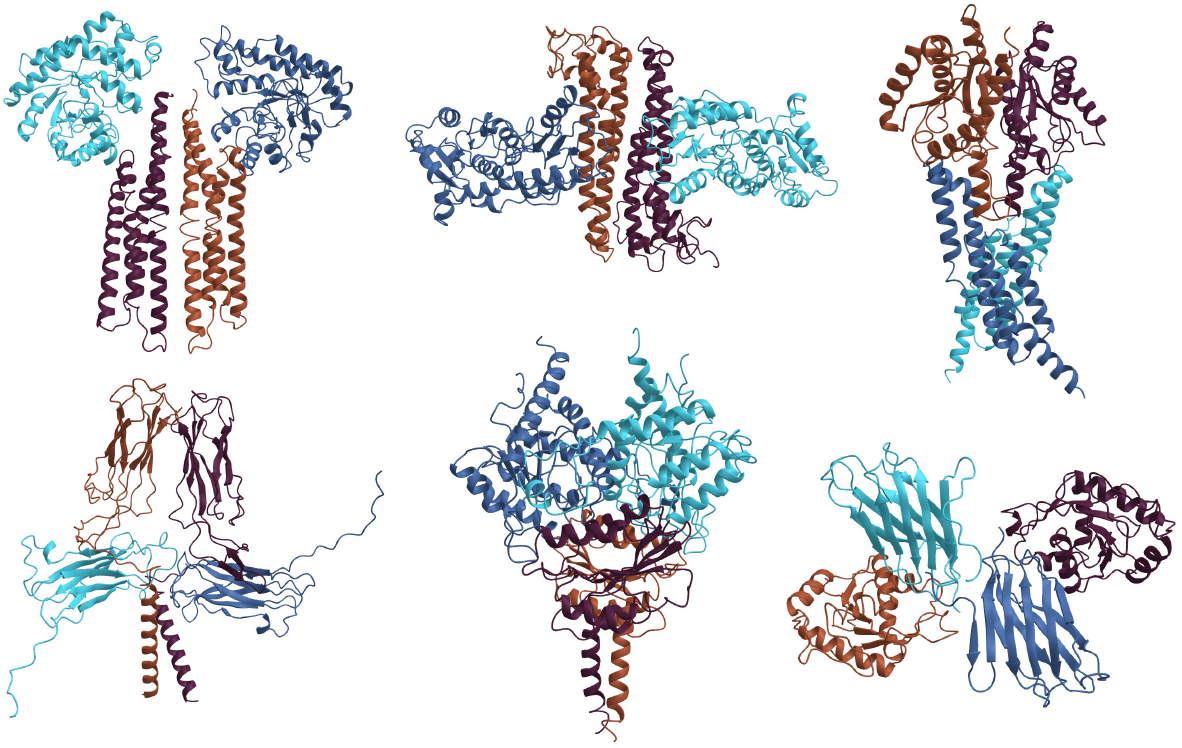
Generations from the trained model, cherry-picked for instances of visually apparent dimer-of-heterodimer structure.

### Symmetry is controlled by a single attention head

ChainStorm has 18 IPA layers (3 × 6), each with 12 attention heads. An investigation of all 216 attention heads suggested that just a single head, the fourth head of the fifth layer of the current frame self-attention IPA (i.e. third from bottom in Figure 1) attends to symmetry, under certain conditions. We will refer to this head as “L5H4” throughout.

We selected four protein complexes from the training set: a homodimer (PDB: 5EDX), a heterodimer where the chains are similar length (PDB: 6R6M), a trimer (PDB: 1TNF), and a monomeric ankyrin repeat, which has a single chain, but a structural motif that repeats throughout the majority of the chain. With these, we constructed (*X*_*t*_, *X*_1_) training pairs for different values of *t*, and visualize the attention weights for L5H4 across *t* for each selected structure.

Figure 4 shows a clear pattern. At *t* = 0, when there is no structural information about the *X*_1_ structure given the information in *X*_0_, L5H4 fires in a diffuse pattern approximately around where the identical residue would be positioned in the symmetric copy of the chain.

**Figure 4:**
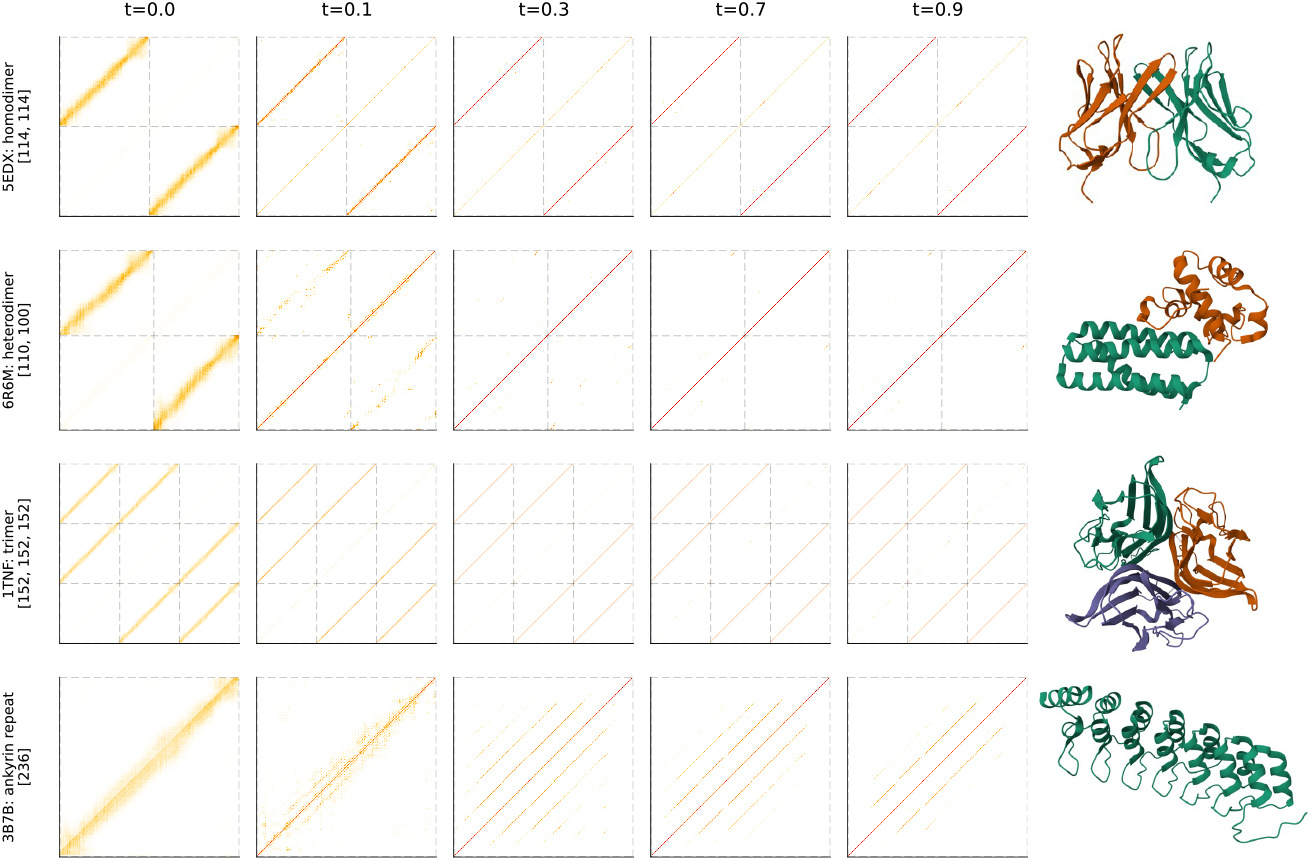
“L5H4”, the 4th head of the 5th *F* self-attention IPA layer (third from bottom in Figure 1) responds to repeating structure. Each heatmap depicts the attention weights, with dashed lines showing chain boundaries, for a training pair *t, X*_*t*_ constructed with *X*_1_ as the PDB depicted. Initially, L5H4 responds with diffuse activation around regions of potential symmetry, which are either concentrated over time, or lost, in which case the head attends to its own amino acid (i.e. along the main diagonal).

Thus for homodimers and length-matched heterodimers the pattern is identical at *t* = 0, concentrated along the two diagonals where the betweenchain residue numbers are similar, with very little activation along the primary diagonal. Note that the relative residue numbering is only provided to the IPA layers within chains, and never between chains, so the network doesn’t have direct access to this signal and must be inferring it.

The same pattern holds for trimers, where corresponding residues in repeating chains have high activation, and in the monomeric ankyrin repeat, at *t* = 0 activation is distribution around the main diagonal.

At *t* = 0.1, the L5H4 activations are less diffuse, having concentrated more precisely on symmetry partner residues, and differences between the homodimers and heterodimers are clearly apparent, where the heterodimer has the majority of activation along the primary diagonal (i.e. no longer attending between chains), which is further reinforced for *t* = 0.3 onwards.

The attention pattern in the monomeric ankyrin repeat shows that L5H4 does not only attend to copies of chains, but also attends to repeating structural motifs within the same chain. The pattern appears to clarify later in the process, and is only clearly visible at *t* = 0.3.

### Suppressing L5H4 prevents symmetry formation

Since L5H4 was the only attention head that appeared to respond to structural similarity or symmetry in training pairs, we hypothesized that we could manipulate its activations to control the generation of symmetric structures.

Figure 5 shows representative samples of dimers generated when input chain lengths are matched. In the absence of any manipulation (left two columns), the dimer complexes are at least approximately symmetric, and the chains comprising the dimer superimpose well upon each other. Also shown (right two columns) is the result of applying a head-specific attention mask to L5H4 that allows it to attend only to the main diagonal (i.e. each residue only attends to itself), which would prevent any patterns that would correspond to symmetry or repeating structural motifs as observed in Figure 4. The generated structures appear visually to be of similar quality to the symmetric dimers, but the superposition clarifies that the structural similarity between the chains is ablated.

**Figure 5:**
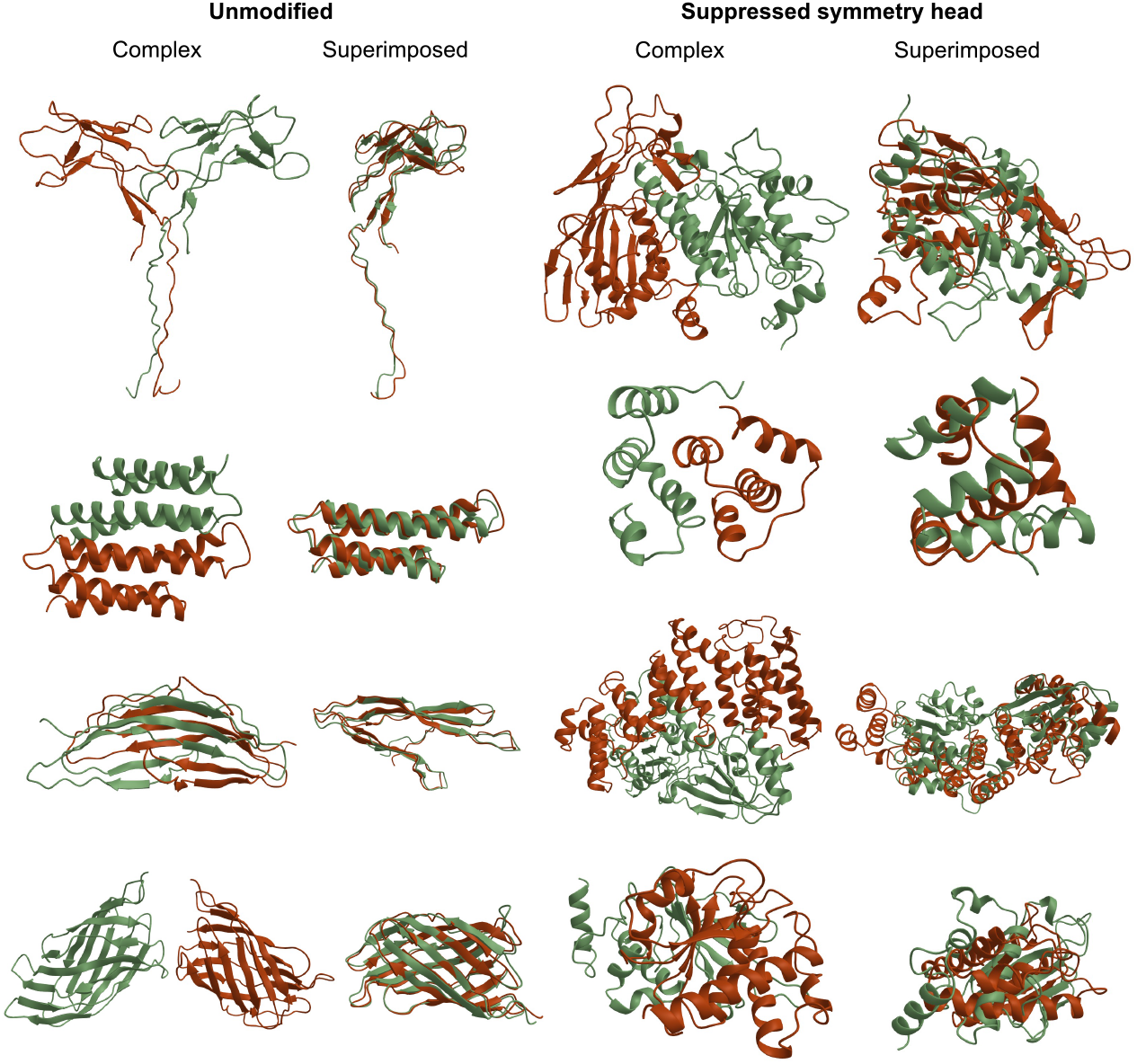
Symmetry head suppression. When initialized with chains of the same length, ChainStorm reliably generates homodimers. The left two columns structures as generated, and the same structures where the two chains have been superimposed, where the similarity between the chains is evident. When the L5H4 attention head is masked so that it can only attend along the main diagonal, the model generates heterodimers (right two columns), evident from the poor match in superimposed structures.

To clarify these observations quantitatively, we require a way to judge multi-chain backbone quality. Self-consistency Template Modeling (TM) scores (scTM) [12] are often used to assess single-chain designed backbone qualities, which rely on generating sequences conditioned on backbones, often using ProteinMPNN [13], and then refolding these *in silico*. However, these approaches are poor at recovering multi-chain structures. Thus to assess the structural quality of the designs, which now involve dimers, we developed a “pseudo multiple sequence alignment” scTM (pMSA scTM) score. As described in Methods, from the designed backbones we first generate 100 sequences (using ProteinMPNN), and then use Boltz-1 [14] to predict the backbone structure using the designed sequences as the MSA input to Boltz1.

With this, Figure 6 shows that suppressing the L5H4 attention head dramatically affects the RMSD between superimposed chains, but does not affect the overall backbone quality when assessed via pMSA scTM score.

**Figure 6:**
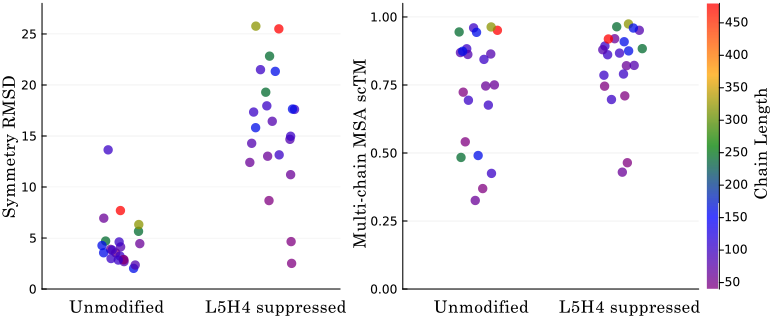
Dimer complexes were generated using ChainStorm, with identical chain lengths. The left panel shows the minimum RMSD for the two designed chains superimposed, where low values indicate an approximate homodimer. The right panel shows the selfconsistency TM score (scTM), here scoring the “refolding” accuracy of the dimer complex (i.e. incorrect binding interaction is penalized), using Boltz1 and re folding from a pseudo multiple sequence alignment of 100 ProteinMPNN sequences designed conditional on the ChainStorm backbones. Each panel compares two cases: “unmodified”, where the standard model is used, and “L5H4 suppressed”, where the single head that appears to control symmetry is prevented from attending between residues of different chains.

### Designed symmetry is driven mainly by chain length similarity and determined early during design

We hypothesized that the primary signal driving the emergence of symmetry was matching chain lengths. To investigate this, we constructed a range of designs where chain A was a fixed length (*L* = 150), and chain B was titrated from 50 to 250. We quantify the “symmetry score” as the proportion of between-chain L5H4 activation (vs. total activation).

Figure 7, Panel A shows that, at *t* = 0 where *X*_*t*=0_ is non-informative and only the chain length information is relevant to the model, the symmetry score varies smoothly with titrated chain B length, and peaks near 1 when the chains A and B are similar in length.

**Figure 7:**
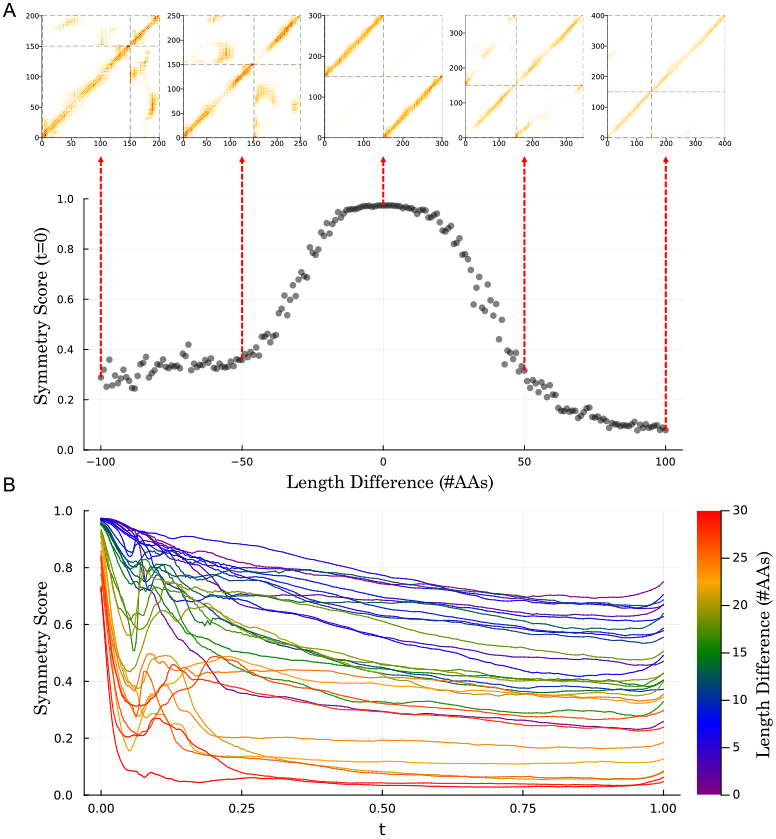
Symmetry trajectories. The 4th attention head of the 5th layer (L5H4) captures whether or not the generated structure will be symmetric, or partially symmetric (with the structure matching for some, but not all, regions). Panel A shows how, at *t* = 0, this head is sensitive to the similarity in the chain lengths. The generation process is initialized with two chains, where one is length 150, and the other varies from 50 to 250, and the L5H4 activations are captures at *t* = 0. The “Symmetry Score” (which is the sum of between-chain activations in this head) is maximal when the chains are of equal length. Heatmaps of head’s activa tion is depicted for 150, 50, 150, 100, 150, 150, 150, 200, and 150, 250 are shown above. Panel B shows how the Symmetry Score evolves over time, where there is a region of high volatility early on in the trajectory where the eventual symmetry is “decided” by the model. Note that the final degree of symmetry is correlated, but not perfectly so, with the difference in chain length.

Titrating Figure 7, Panel B investigates the full generative timeline for chains of similar length, titrating chain B between 150 and 170 AAs. At *t* = 0 the L5H4 all begin near 1, but are greater for more similar chain lengths, as expected from Panel A. But as the structures are generated, the symmetry score declines rapidly for chains of greater length difference, and then experiences a strong fluctuation between *t* = 0.05 and *t* = 0.15, and then is relatively stable for the rest of the process.

Note that, from this sampling, some chains of very similar length, including one with *L*_*A*_ = 150, *L*_*B*_ = 151 terminated with very low symmetry scores, indicating that the model is capable of spontaneously designing heterodimers of similar chain lengths.

## Discussion

We demonstrate that a relatively simple generative model of protein structure and sequence learns to spontaneously generate multi-chain protein complexes that, even in the absence of any symmetry constraints during generation, exhibit patterns of symmetry that are observed in the PDB training data. We have isolated this behavior to a single attention head, which activates for pairs of residues that match across symmetric chains, or that match across repeating structural motifs even within the same chain, and confirmed that suppressing these activations prevents the model from designing symmetric chains.

Existing models are capable of generating symmetric structures when some form of symmetry constraint is applied. To generate an oligomer, for example, RFdiffusion [15] initializes a monomeric initial state that is then copied by an explicit symmetry operation, which is enforced during the generative process. Also, models of protein design that manipulate sequence space (relying on a folding model to generate a backbone conditioned on the sequence) can also capably design symmetric assemblies when a symmetry constraint is applied to the sequences [16].

ChainStorm is, to the best of our knowledge, the first model to spontaneously generate symmetry in the absence of any symmetric initialization or symmetry constraints applied during generation. It is not surprising that such patterns can be learned, given how common they are in the training data. One question then is why this is not observed in other models. A survey of models suggests that the overwhelming majority of them are trained on single-chain structures, either entirely or where a multi-chain training phase is finetuning only, which may not be sufficient to induce this behavior. A notable exception is Chroma [17], which is trained on multi-chain structures. But given the nature of how ChainStorm identifies symmetry, we hypothesize that Chroma’s Graph Neural Network approach permits only sparse random connectivity between distal pair-wise residues, which may hinder the model’s ability to perceive patterns of symmetry between chains.

ChainStorm’s single attention head that attends to structural repetition is reminiscent of the “induction head” phenomenon in autoregressive large language models of text, where a class of attention heads specializing in identifying repetition [18], [19]. In the autoregressive LLM context, induction heads identify past instances of any text motifs that the current token might echo, and act to increase the probability of the tokens that had previously succeeded the current one. In LLMs, these induction heads are consequential. Their emergence, and the LLM “learning to copy”, corresponds to a noticeable drop in the model’s loss, and it has been argued that they are the primary circuit responsible for “in context learning” exhibited by LLMs.

Our context differs because our model is not autoregressive, and we are not aware of any work on induction head-like circuits in text diffusion models. Further, instead of tokens we have rotational frames. Nevertheless, the L5H4 head is clearly driving the model’s ability to be able to copy structure, both between and within chains.

### Future Directions

Many protein design tasks involve some sort of symmetry. This includes the de novo design of multimers, including dimers, trimers, etc, as well as large self-assembling icosahedral multimers [20], but it also includes tasks where components of an existing multimer need to be redesigned while maintaining symmetry. One avenue for future work will explore whether a model, like ChainStorm, that is symmetry-aware would have some sort of advantage over a model that is not natively symmetry aware [15], when design is performed under symmetry constraints, as the model’s predicted *t* = 1 state might be more aligned with the symmetry constraints.

By inspecting the flow trajectories during inference, we note that there is a sudden transition where the predicted target to which the process is flowing suddenly appears, often within a very short time window. As can be seen in Figure 2, this also varies by chain in a way where e.g. common domains (like the immunoglobulin heavy and light chains in the figure) resolve earlier and more precipitously than other less common chains. We hypothesize that, since flow matching learns to match the density of the target distribution, common chains will have very wide basins of attraction, and when the current state is within that basin the *t* = 1 endpoint is, macroscopically at least, fully determined, since the process noise cannot cause the state to exit that basin due to its size. Sudden transitions in the predicted *t* = 1 state may cause high curvature in the sample paths, and inference might be improved if this could be avoided. Further, with self-conditioning, the order in which events resolve may be critical for the model’s ability to learn to build complexes. We anticipate that modifying the process may reduce these effects, but it will require detailed empirical investigation to understand whether this is beneficial or not.

Finally, this model is, in a sense, a “base” model, and is not currently useful for any specific task besides unconditional structure generation. Future work will involve building in various conditioning strategies, allowing the model to sample binders to targets, to redesign parts of a chain, or to scaffold motifs. Finally, ChainStorm makes no attempt to sample from the restricted space of chains that can be predicted after inverse-folding sequence generation, which has shown to be useful for in silico design screening. We intend to explore strategies to allow this, without damaging the models ability to sample from diverse symmetric structures.

### Limitations

While our model co-designs sequence and structure, we have not investigated the quality of the designed sequences at all. In many sequence/structure co-design models the sequence quality is not as good, for e.g. in silico refolding [21], as sampling a sequence conditioned on the backbone [13], and we do not expect ChainStorm to be an exception here.

We also do not explore which components of the model are critical for inducing, during training, the spontaneous learning of symmetry, nor do we explore whether the result that the symmetry is primarily encoded by a single attention head in a single layer is a chance event, or a reliable outcome.

## Methods

### Notation and state space

Let *L* be the number of residues.

The designable component of a protein backbone and sequence is represented on the product manifold

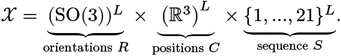

A generic protein backbone is

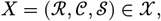

with

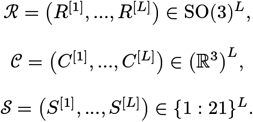

A noise-corrupted sample will be indexed with time: *X* = (ℛ _*t*_, 𝒞 _*t*_, 𝒮 _*t*_) each with elements 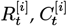 and 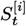.

### Model

The model, ℳ takes the time *t*, the current backbone “frames” *F* = (ℛ_*t*_, 𝒞_*t*_), and the current Amino Acids (one-hot encoded) 𝒮_*t*_, and a set of Pair Features that encode the distance between two residues in the primary sequence, and whether or not two residues are part of the same chain. The model returns an estimate of the frames, 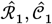 and the logits of the amino acid sequence 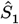. The model uses “self-conditioning”, and additionally takes *F*_SC_ = (ℛ_SC_, 𝒞_SC_), which are either the predicted backbone frames from a previous pass of the model, or nothing:

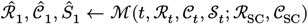

The model iteratively transforms *D* × *L* embeddings (*D* = 384 in this version of the model). These are initially projected from the amino acids, and updated by the model “trunk” which repeats 6 times. In each repeat, the self-conditioning frames *F*_SC_ first interact with each other via Invariant Point Attention (IPA), then *F*_SC_ interacts with *F* via a cross-attention variant of IPA, and then frames in *F* interact with each other via IPA.

The pair features are encoded via Random Fourier Features (RFF) and fed into each IPA layer, and diffusion time *t* is encoded via RFF and fed to an adaptive layernorm (AdaLN) prior to each IPA layer, as well as prior to the SwiGLU feed-forward layer. In each iteration of the trunk, the frames *F* are updated via a function of a linear projection of the embeddings (as in AlphaFold2). The model outputs the final state of *F* and the amino acid logits which are projected from a SwiGLU fee-forward layer after completion of the trunk. See Figure 1 for a depiction.

### Pair features

Between each pair of amino acids *i* and *j*, with residue numbers *n*_*j*_, *n*_*j*_, and chain indices *c*_*i*_ and *c*_*j*_, we encode two features: s

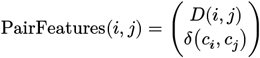

where *δ*(*c*_*i*_, *c*_*j*_) = 1 if *c*_*i*_ = *c*_*j*_, else 0, and where

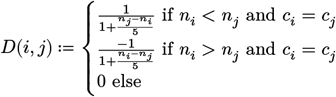

Thus for any pair of positions, the model is informed of i) whether or not they are from the same chain, ii) whether or not one is ahead of the other in the primary sequence (for residues of the same chain) via the sign of *D*(*i, j*), and iii) the distance (i.e. number of amino acids) between them in the primary sequence, which is scaled to change rapidly for nearby neighbors and slowly for distant ones. Figure 8 shows *D*(*i, j*) for residues from the same chain.

**Figure 8:**
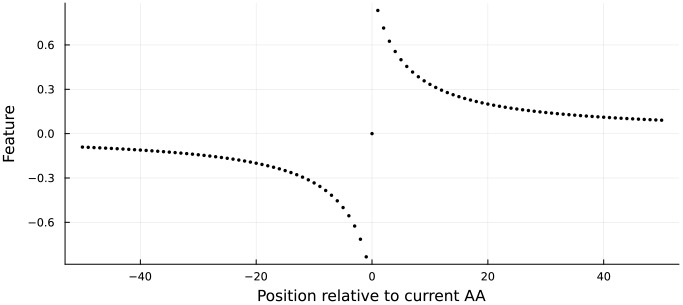
Encoding of positional residue differences.

### Endpoint-conditioned sampling

It is convenient to describe our training and inference process for these states in terms of Step functions that take a guided stochastic step from *X*_*t*_ to *X*_*t*+Δ*t*_, conditioned on terminating at an endpoint *X*_∗_ (either an *X*_1_ observation or an 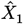 prediction at *t* = 1). For each state space we also define a Bridge function, which draws a state at *t* conditioned on states at *t* = 0 and at *t* = 1.

We describe the action of Step and Bridge on single elements *R*_*t*_, *C*_*t*_, and *S*_*t*_ (omitting the superscript positional index), and their action on a state ℛ_*t*_, 𝒞_*t*_, and 𝒮_*t*_ is upon all elements of the state, independently.

### Orientation

With noise scale *η*, for one orientation *R*_*t*_ with endpoint *R*_∗_:

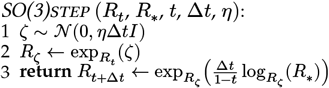

where *ζ* is sampled in the tangent space of *R*_*t*_. *R*_*t*_ is thus noised with variance proportional to *η*Δ*t*, and then moved towards *R*_∗_. To sample *R*_*t*_ ∼ Bridge (*R*_0_, *R*_1_, *t*) when 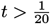, multiple smaller SO(3)steps, each with size 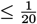, are taken through intervening time points until *t* is reached.

### Position

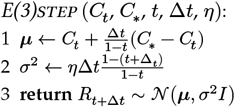

This is the exact solution and for any 0 < *t* < 1, *C*_*t*_ ∼ Bridge (*C*_0_, *C*_1_, *t*) can be sampled with a single E(3)step.

### Sequence

Following equation 10 from [8], which constructs the distribution at *t* as a mixture of *S*_1_, uniform noise (i.e. all states are equally likely), and *S*_0_, where the mixture weights vary over time according to specified “noise schedule” functions. Here we use 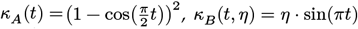, and *κ*_C_ (*t*) = (1 *−* (*κ*^[1]^ + *κ*^[2]^)). With *S*_0_, *S*_1_ as a one-hot probability vectors:

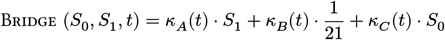

Applying Theorem 3 to equation 10 in [8], we obtain:

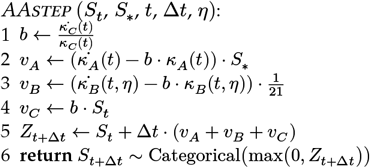

Our initial *S*_0_ state is all “dummy character”, with *S*_0_ = 21. Note that the “uniform noise” *K*_*B*_(*t*) mixture component implies that, during training, some amino acids will match neither *S*_0_ nor *S*_1_. During inference, if the model “unmasks” a character early that later turns out to not be compatible with the backbone or the other characters, these unmasked characters to switch again.

### Schedules

For *R* and *C* the diffusion scale 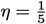 and, aiming to have the amino acids resolve later than the backbone, for *C* and *R* (but not *S*) we rescale time with *τ* ≔ *φ*(*t*), where

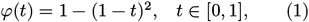

The schedule for *S* is incorporated directly into the AAstep procedure. The relative timing and stochasticity of these Step procedures is visualized in Figure 9

**Figure 9:**
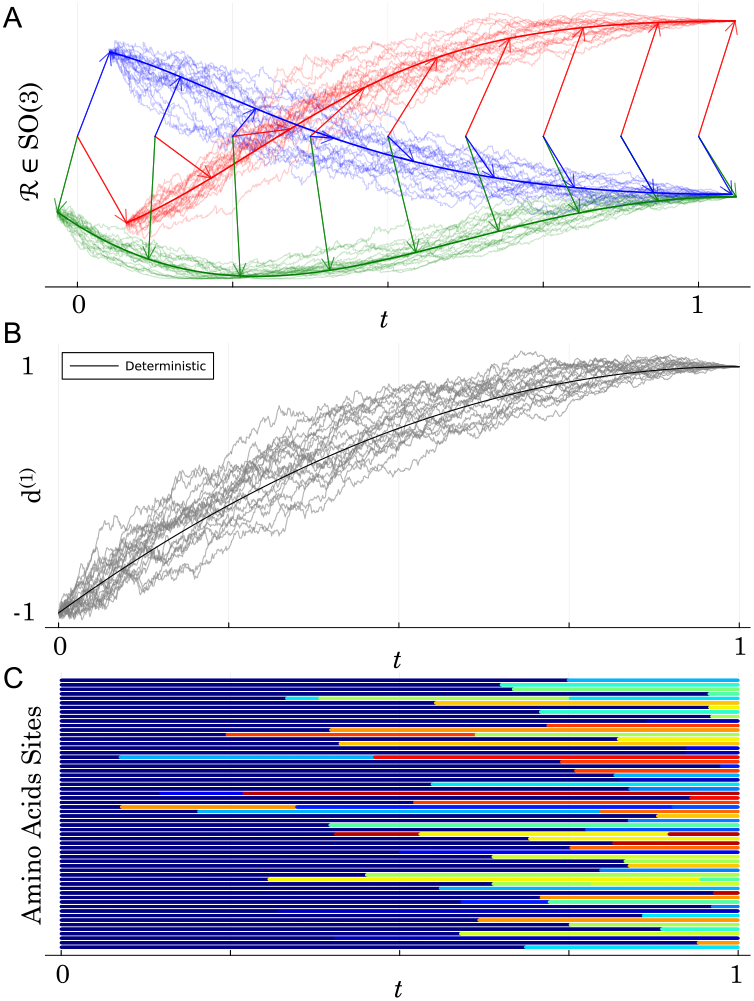
Stochastic interpolation. ChainStorm’s training and inference rely on stochastic interpolation, where a noisy process is sampled conditioned on two endpoints. Panel A shows the diffusion for rotations, using a SO(3) manifold bridge between two endpoints, with 20 noisy realizations (faint lines) as well as the deterministic (i.e. timer-escaled geodesic) path. Panel B depicts a single coordinate position (i.e. one of the dimensions of the centroid of a backbone frame) under a timerescaled Brownian bridge, if its value was −1 at *t* = 0 and 1 at *t* = 1. Also depicted are 20 replicates of the noisy bridge as well as the deterministic trajectory (i.e. if the Brownian motion noise was 0), depicted in bold. Each horizontal line in panel C shows a realization of the discrete process that interpolates between characters, starting at *t* = 0 with a dummy 21st state, and switching stochastically, with some trajectories having multiple switches.

### Training

From the training distribution we draw instances of *X*_1_ = (*R*_1_, *C*_1_, *S*_1_), a (possibly multi-chain) “biological assembly”, and instances of *X*_0_ = (*R*_0_, *C*_0_, *S*_0_) ∼ *p*, with 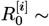 Uniform(SO(3)), 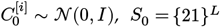.

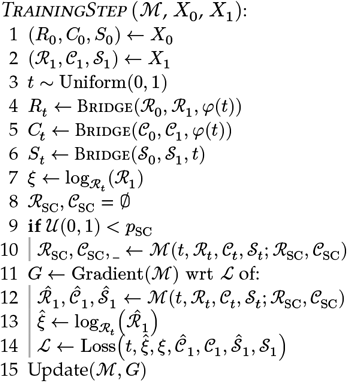

Computationally, *ξ* and 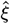 are represented as coordinates in the canonical orthonormal basis. *p*_SC_ controls the proportion of training steps that use self-conditioning.

### Time-dependent loss scaling

The loss can be scaled as a function of time to stabilize training. For example, the standard Conditional Flow Matching (CFM) objective in Euclidean space takes a uniformly weighted expectation over time

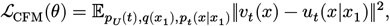

where *p*_*U*(*t*)_ = 1 for any 0 < *t* < 1 and 0 elsewhere, corresponding to *t* ∼ Unif(0, 1), and where the model *v*_*t*_(*x*) predicts 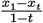 for each training pair:

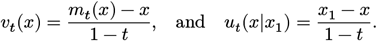

Here instead, our loss is over *x*_1_ directly, with

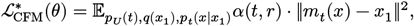

compensating for this reparametrization in the loss by explicitly scaling it by *α*(*t, r*):

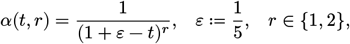

where *r* = 2 for the position and orientation components, *r* = 1 for discrete components, and *ε* is chosen to stabilize the loss across *t*. Note this does materially change the loss. In Euclidean space, the loss up to a constant *K*_*r*_ becomes:

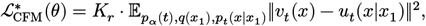

where, instead of *t* being uniformly weighted in the expectation, it is weighted by:

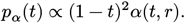

Indeed, since

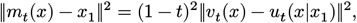

we can rewrite 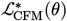 as

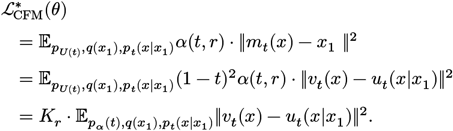

Here, *K*_*r*_ specifies the normalizing constant for the density *p*_*α*_ ∝ (1 *− t*)^2^*α*(*t, r*), via

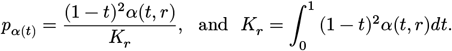

Therefore, the loss 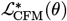, up to a constant, coincides with the CFM objective taking the expectation over *t* ∼ *p*_*α*_ instead of *t* ∼ Unif(0, 1).

For example, when taking the loss over position we will have *r* = 2, and thus

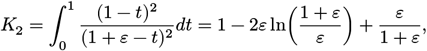

Where

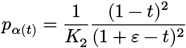

specifies the reparametrized distribution *t* ∼ *p*_*α*_.

For a sufficiently flexible model this should not matter, but for a finite model this allows control over the relative importance of features that emerge in the conditional paths at different *t*. Compared to the standard uniform weighting (where the implied loss scaling spikes as *t* → 1 to compensate for the expected difference between *x*_*t*_ and *x*_1_ approaching 0) our choice progressively suppresses the finer details as *t* → 1, which is reasonable in this context as they will be largely driven by meaningless atomic noise in the training data. Figure 10 shows how this choice re-weights the expectation over *t* in the 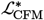 loss.

**Figure 10:**
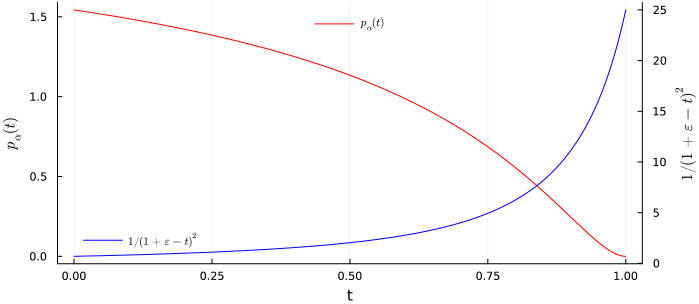
Our loss scaling, with 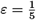, for the position and orientation components. Shown are the actual loss multiplier, in red, and the distribution this induces over the Conditional Flow Matching expectation over *t*.

### Loss

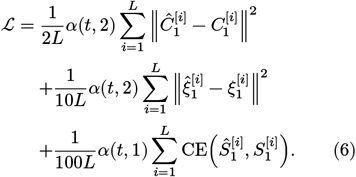

with CE the cross-entropy against the one-hot training amino acid sequence, and 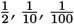 aim to balance the loss of the three components. Note that we use a different power in the CE loss term: *α*(*t*, 1).

### Sampling

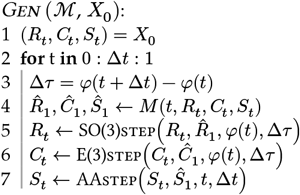

### Data setup

We train ChainStorm on a set of protein structures from the Protein Data Bank (PDB) [11] with entries deposited up until February 2025. For each entry, we use BioStructures.jl [22] to download the mmCIF file of the first biological assembly, filtering out all non-peptide chains and molecules. For consistent and strictly increasing residue numbers without insertion codes, while still respecting gaps, we renumber using unique and sequential residue numbers found at `_atom_site.label_seq_id` in the mmCIF format.

We primarily compiled this dataset for our other models, which make use of some of the additional metadata present in the asymmetric unit versions of the mmCIF files. For this reason, we only include PDB entries where we were able to load both the biological assembly and asymmetric unit versions using BioStructures.jl v4.

To acquire the frame of residue *i*, we run the Kabsch algorithm [23] to align the backbone atom coordinates *N*^[*i*]^, *Cα*^[*i*]^, and *C*^[*i*]^ to backbone atom coordinates of a centered and idealized template residue

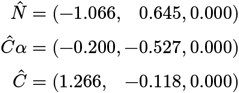

from which we obtain a rotation matrix *R*^[*i*]^ and translation vector (centroid) *C*^[*i*]^ such that each atom coordinate ***a*** in the template frame is transformed to the residue frame as *R*^[*i*]^***a*** + *C*^[*i*]^.

We address overrepresentation of certain protein families in the PDB by assigning a “cluster” label to each structure, defined as the index of connected component on a heuristic sequence similarity graph. This prevents the model from being dominated by abundant proteins.

The preprocessed dataset is stored at huggingface.co/datasets/MurrellLab/ProteinChains/ and can be loaded with DLProteinFormats.jl

### Training

The model was trained with a maximum record length of 1500 amino acids, batching together records of similar length, with a batch dimension of 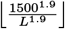. Since the training self-conditioning pass occurs outside of the gradient calculation, the model was first trained for five epochs without self-conditioning (i.e. *p*_SC_ = 0 in TrainingStep), to ensure the self-conditioning predictions would be reasonable when first encountered. The two self-conditioning IPA layer outputs were set to 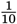 of their default initial magnitudes to minimize the initial effect of untrained self-conditioning layers, and then the model was trained for either 10 epochs or 28 epochs, followed by a two-epoch linear learning rate warmdown in both cases. Results described in this manuscript were from the 28 additional epoch model (“checkpoint 31”), but the emergence of symmetry was clearly evident in the earlier checkpoint as well. All training was performed on a single NVIDIA RTX 6000 Ada.

### pMSA scTM

We propose a refolding pipeline for multimeric backbone complexes. For a given protein, we generate *N* sequences using a sequence-given-structure model. Then we use a folding model which takes one sequence to fold as input, but also an optional Multiple Sequence Alignment (MSA). We pass in the first of the N sequences as the sequence to be folded and then we pass in all the N sequences as the MSA input. We then define the pseudo MSA scTM score (pMSA scTM) as the TM score between the original generated structure and the output structure from the folding model.

We used ProteinMPNN [13] as the sequence-given-structure model and Boltz1x [14] as the folding model, and set *N* = 100.

The impact of the pMSA seems to be especially high for protein complexes, as shown in Figure 11.

**Figure 11:**
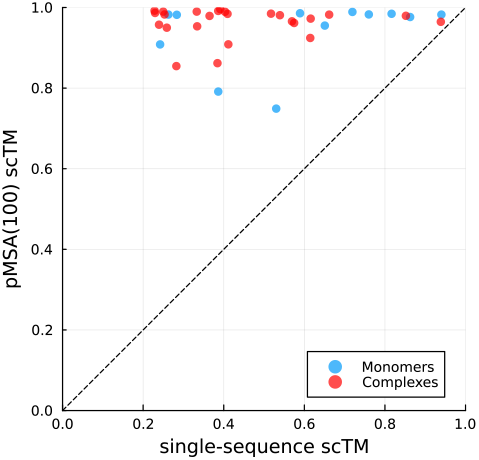
Single-sequence scTM and pMSA scTM with *N* = 100 computed for structures from the publicly available CASP15 targets [24], removing structures with large non-protein ligands. Since many structures had missing residues, we only generated the sequence for the available residues and filled in the missing gaps with the original sequence. Only the sequence to be folded had gaps filled in, with the pMSA keeping the gaps.

**Figure 12:**
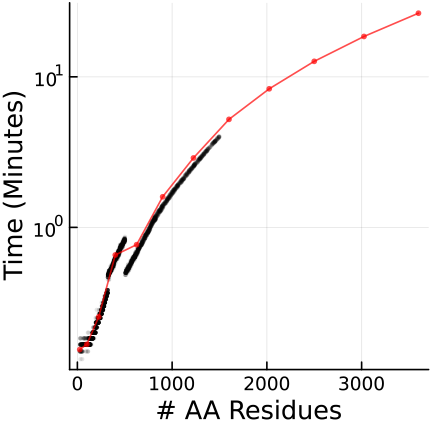
Sampling speed, in minutes, using a single NVIDIA RTX6000 Ada GPU, and default sampling parameters. Black shows timing between 25 and 1500 residues for the samples from the UMAP plot, showing that the timing is very consistent for a given length, and red (run on a different machine with the same GPU model, but a different CPU) extends these with single points out to 3600 residues (which took 26 minutes).

### Implementation and availability

ChainStorm is implemented entirely in Julia [25], a language for Scientific Computing. From the general Julia package ecosystem, we use Flux.jl [26] as a deep learning framework, Zygote.jl [27] for automatic differentiation, and CUDA.jl [28] for GPU support. Makie.jl [29] and Plots.jl [30] were used for visualization and plotting, and Biostructures.jl [22] was used for PDB imports and preprocessing.

ChainStorm itself is available at:

- github.com/MurrellGroup/ChainStorm.jl

Much of the functionality behind ChainStorm comes via a number of open source mutually interoperable Julia packages developed by us, which have mostly not been previously described in manuscripts, and are available under github.com/MurrellGroup:

- `Flowfusion.jl`
  ▸ Generic flow matching/diffusion implementation for euclidean, manifold, and discrete spaces.

- `ForwardBackward.jl`
  ▸ Implementation of some stochastic processes used by `Flowfusion.jl`

- `InvariantPointAttention.jl`
  ▸ For the core IPA layers (previously used in [4])

- `BatchedTransformations.jl`
  ▸ For GPU-differentiable batch-dimension-augmented affine transforms, including rigid trans formations.

- `Onion.jl`
  ▸ For assorted layers, including Adaptive Layernorm, SwiGLU, Framemover, and some InvariantPointAttention.j/ BatchedTransformations.jl interoperability.

- `CannotWaitForTheseOptimisers.jl`
  ▸ Implements a number of bleeding-edge optimizers, including Muon (originallyimple mented here: [31]) which was used to train ChainStorm

- `ProteinChains.jl`
  ▸ Representations and transformations of protein structure data

- `ProteinRefolding.jl`
  ▸ Used here for pMSA scTM score calculations.

- `ProtPlot.jl`
  ▸ For backbone geometry visualization (via Makie.jl)

- `DLProteinFormats.jl`
  ▸ For preprocessed GPU/DL compatible tensorformatted PDB data.

## Acknowledgements

This project was funded by the Swedish Research Council (2023-02516) to B.M.

